# Deep Learning with Multimodal Representation for Pancancer Prognosis Prediction

**DOI:** 10.1101/577197

**Authors:** Anika Cheerla, Olivier Gevaert

## Abstract

Estimating the future course of cancer is invaluable to physicians; however, current clinical methods fail to effectively use the vast amount of multimodal data that is available for cancer patients.

To tackle this problem, we constructed a deep neural network based model to predict the survival of patients for 20 different cancer types using gene expressions, microRNA data, clinical data and histopathology whole slide images (WSIs). We developed an unsupervised encoder to compress these four data modalities into a single feature vector for each patient, handling missing data through a resilient, multimodal dropout method. Encoding methods were tailored to each data type - using deep highway networks to extract features from genomic and clinical data, and convolutional neural networks extract features from pathology images. We then used these feature encodings trained on pancancer data to predict pancancer and single cancer survival data, achieving a C-index of 0.784 overall.

This work shows that it is possible to build a pancancer model for prognosis that also predicts prognosis in single cancer sites. Furthermore, our model handles multiple data modalities, efficiently analyzes WSIs, and summarizes patient details flexibly into an unsupervised, informative profile. We thus present a powerful automated tool to accurately determine prognosis, a key step towards personalized treatment for cancer patients.

## Introduction

Estimating tumor progression or predicting prognosis can aid physicians significantly in making decisions about care and treatment of cancer patients. To determine the prognosis of these patients, physicians can leverage several types of data including clinical data, genomic profiling, histology slide images, and radiographic images, depending on the tissue site. Yet, the high-dimensional nature of some of these data modalities necessarily implies that physicians can not always manually interpret these multimodal biomedical data to determine treatment and estimate prognosis [1, 2].

The presence of inter-patient heterogeneity warrants that characterizing tumors individually is essential to improving the treatment process [3]. Previous research has shown how molecular signatures such as gene expression patterns can be mined using machine learning and are predictive of treatment outcomes and prognosis. Similarly, recent work has shown that quantitative analysis of histopathology images using computer vision algorithms can provide additional information on top of what can be discerned by pathologists [4]. Thus, automated machine-learning systems, which can discern patterns among high-dimensional data may be the key to better estimate disease aggressiveness and patient outcomes. Another implication of inter-patient heterogeneity is that tumors of different cancer types may share underlying similarities. Thus, pancancer analysis of large-scale data across a broad range of cancers has the potential to improve disease modeling by exploiting these pancancer similarities.

Multi-institutional projects such as The Cancer Genome Atlas (TCGA) [5–7], which collected standardized clinical, multi-omic and imaging data for a wide array of cancers, are crucial to enable this kind of pancancer modeling.

Automated prognosis prediction however remains a difficult task mainly due to the heterogeneity and high dimensionality of the available data. For example, each patient in the TCGA database has thousands of genomic features (e.g. microRNA or mRNA) and high resolution histopathology whole slide images (WSIs). Yet, based on previous work, only a subset of the genomic image features are relevant for predicting prognosis. Thus, to successfully develop a multi-modal model for prognosis prediction, an approach is required that can efficiently work with clinical, genomic and image data, in essence multimodal data. Here, we tackle this challenging problem by developing a pancancer deep learning architecture drawing from unsupervised and representation learning techniques, and developing a resilient learning architecture that exploits large-scale genomic and image data to the fullest extent.

The main goal of this contribution is to harness the vast amount of TCGA data available to develop a robust representation of tumor characteristics that can be used to cluster and compare patients across a variety of different metrics. Using unsupervised representation techniques, we develop pancancer survival models for cancer patients using multi-modal data including clinical, genomic and WSI data.

### Background

Prognosis prediction can be formulated as a censored survival analysis problem [8, 9], predicting both *if* and *when* an event (i.e. patient death) occurs within a given time period. Given the unique statistical distribution of survival times, they are canonically parametrized using the “hazard function”, such as in standard Cox regression.

In recent years, many different approaches have been attempted to predict cancer prognosis using genomic data. For example, Zhang *et al.*, used an augmented Cox regression on TCGA gene expression data to get a *C*-index of 0.725 in predicting glioblastoma [10]. MicroRNA data in particular have shown high relevance as a measure for disease modeling and prognosis [11–14], with Christinat *et al.*, achieving a *C-index* of 0.77 on a subset of renal cancer data using random forest classifiers [15]. However, despite the high performance of machine learning models based one molecular data alone, there is still some scope for improvement; after all, the tumor environment is a complex, rapidly evolving milieu [16–18] that is difficult to characterize through molecular profiling alone.

Recently, the use of whole slide image (WSI) data has been shown to improve the performance and generality of prognosis prediction. As WSIs are high resolution images of cellular architecture and environment with potentially only a fraction of the slide relevant to predicting prognosis, much of the literature focuses on hybrid approaches involving pathologist annotation of regions of interest (ROIs). For example, Wang *et al.*, match the performance of genomic models by using 500 by 500 pixel, physician-selected ROIs and handcrafted slide features to predict prognosis [19]. More recently, deep learning provides a significant boost in predictive power. For example, Yao *et al.*, are able to significantly outperform all molecular profiling-based methods on two lung cancer data sets using only physician-selected ROIs and convolutional neural networks (CNNs) [20]. Other reports, including Beck *et al.* and Bejnordi *et al.* showing that histopathology image data contains important prognostic information that is complementary to molecular data [21, 22]. Yet, multimodal prognosis models are still highly underexplored [23]. To our knowledge, only one paper explores combining genomic and image data for prognosis, showing that a lung-cancer genomic model (C-index 0.660) and WSI-based model with hand-annotated ROIs (C-index 0.613) can be combined to get a final classifier with C-index 0.691 [24].

Moreover, the WSI-based methods discussed above require a pathologist to hand-annotate ROIs, a tedious task. Arguably the most difficult part of automated, multimodal prognosis prediction is finding clinically relevant ROIs automatically. In the related field of tumor classification from WSIs, a “decision-fusion” model that randomly samples patches and integrates them into a Gaussian mixture has yielded accurate predictions [25]. Moreover, more recent work has focused on using attention mechanisms to learn what patches are important [26]. However, in prognosis prediction, truly-automated WSI-based systems have had limited success. One report uses a slide-based approach that relies on unsupervised learning – Zhu *et al.*’s recent paper uses K-means clustering to characterize and adaptively sample patches within slide images, achieving 0.708 C-index on lung cancer data [27], a result that nearly rivals genomic-data approaches.

All previous research has focused on only single-cancer data sets, missing the opportunity to explore commonalities and relationships between tumors in different tissues. And although previous papers explore both genomic and imaging based approaches, few models have been developed that integrate both data modalities. By exploiting multimodal data, as well as developing better methods to automate WSI scoring and extract useful information from slides, we have the potential to improve upon the state-of-the-art.

In recent years, CNNs have been used to significantly improve machine learning tasks [28] including missing value estimation in genomic data [29] and prediction of prognostic factors based on WSI [26]. A key component of the success of CNNs is their ability to deal with high-dimensional, unstructured data, in particular image data [30]. For example, CNNs can accurately classify scenes from images by learning a set of flexible, hierarchical features [31]. Even if the majority of pixel inputs are “dropped out” completely for some samples, this model can still be trained to predict accurately and can handle the uncertainty [32].

The prognosis prediction task is more unstructured than traditional deep learning tasks; instead of classifying from relatively small images (224 by 224 for ImageNet, for example), we must predict survival times from biopsy slides that are much larger. Furthermore, patients span a wide variety of cancer types, and are often missing some form of imaging, clinical, or genomic data, making it difficult to apply standard CNNs. Unsupervised learning has shown significant promise [33]. By learning unsupervised correlations among imaging features and genomic features, it may be possible to overcome the paucity of data labels. Similarly, representation learning techniques might allow us to exploit similarities and relationships between data modalities [34]. In prognosis prediction, it is crucial that the model maps similar patients to the same abstract representation in a way that is agnostic to data modality and availability. Taking inspiration from unsupervised and representation learning should help tackle many of the challenges that make prognosis prediction using multimodal data difficult.

## Materials and Methods

### Data Sets and Tools

Our main source of data is the TCGA database, (http://cancergenome.nih.gov/) [5–7], which contains microRNA data for 1,881 microRNAs, gene expression data for 60,383 genes, a wide range of clinical data, and WSI data for over 11,000 patients. Table 1 describes the data distribution in more detail. It is clear that many patients do not have all their data available, implying that classifiers and architectures that can deal with missing data might perform better. Each patient has a time of death recorded, right-censored up to a maximum of 11,000 days after diagnosis across all cancer sites. The 20 cancers we examine have significantly different survival patterns, as can be seen in Fig 1. We rely on the Python package *openslide* to efficiently read and parse WSIs and the *PyTorch* framework to enable the creation of neural network models. To train our models, we use an NVIDIA™ GTX 1070 GPU.

**Table 1.** Data distribution of TCGA data including missing data. Survival data is available for the majority of patients, while microRNA and clinical data are missing in a subset of patients. Nearly 43% of patients have at least one type of missing data.

| Data type | Number of cases | Number of missing cases | Percentage missing |
| --- | --- | --- | --- |
| gene expression data | 10198 | 962 | 8.62% |
| microRNA expression data | 10125 | 1035 | 9.27% |
| WSI slide data | 10914 | 246 | 2.2% |
| clinical data | 7512 | 3648 | 32.69% |
| survival target data (time of death) | 11121 | 39 | 0.35% |
| patients with complete data | 6404 | 4756 | 42.62% |

**Fig 1.**
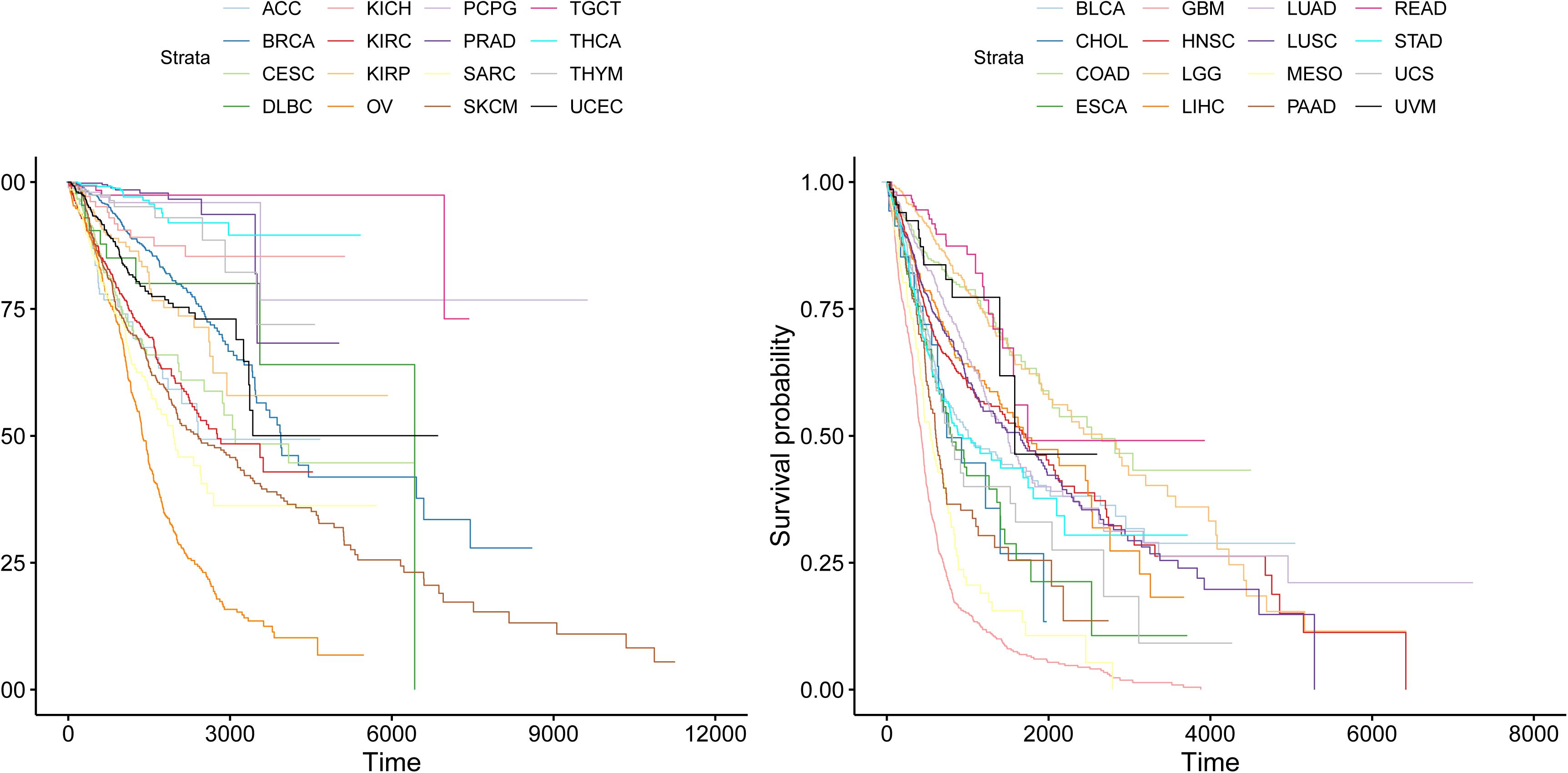
Kaplan-Meier survival curves for all cancer sites in the TCGA database showing that survival is very tissue specific.

The TCGA dataset of 11,160 patients was split into training and testing data sets in 85/15 ratio, stratifying by cancer type in order to ensure the same distribution of cancers in both the training and test sets.

### Deep Unsupervised Representation Learning

In order to train a pancancer model for prognosis prediction, we first attempt to compress multiple data modalities into a single feature vector that represents a patient. Previous work has found significant cross-correlations between different data types (e.g. gene expression, clinical, microRNA and image data) [23, 35], and learning these relations in an unsupervised fashion could significantly improve the prognosis prediction process. Thus, we use a representation learning framework to guide our approach.

Although approaches such as split-brain autoencoders induce convergence between different multimodal feature representations, they rely on reconstruction error, which may not be a good choice for heterogeneous data sources. Instead, we rely on a method inspired by Chopra *et al.*, in which two different “views” of objects are passed through a Siamese network to create feature representations [36]. For views from the same object, the cosine similarity between these feature representations is maximized, whereas for views from different objects, the cosine similarity is minimized. To ensure stability, a margin-based, hinge-loss formulation is used, such that different-object feature representations are only penalized if they fall within a margin *M* of the same-object representations. This forces different views of a single patient’s information to have similar feature vectors, while avoiding mode collapse where all features predict exactly the same vector for all patients.

In this work, we use a similar formulation as [36], but with some modifications. Because of the different data modalities, instead of using a Siamese network, we use one deep neural network for each data type, with differing architectures described in Fig 2. We define the feature space to have a length of 512. Since we have more than two different modalities, we sum over the similarity loss for each pair of modalities that are present. We can define the loss *l*_*sim*_(*θ*) as in the equations 1, 2 and 3.

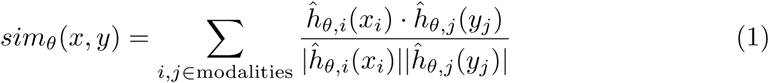

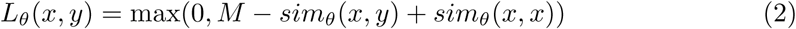

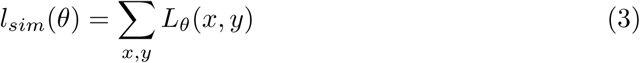

where *x*_*i*_ is the data for modality *i* and *ĥ*_*θ,i*_ is the predictive model for modality *i*. Note that the parameter *M* controls the “tightness” of the clustering. If *M* is high, feature vectors for a given patient are permitted to be relatively different, as long as they stay similar to a certain extent. If *M* is low, feature vectors for a patient are forced to be much closer together, which is usually more ideal, but can also cause mode collapse. After parameter search, we settled on *M* = 0.1 as the default value because it was the smallest value of *M* that didn’t cause mode collapse. This loss is computed between every pair of patients in a batch. Thus, the unsupervised model must learn to recognize important, patient-distinguishing patterns in genomic and image data. Moreover, it must learn how patterns in one modality correspond to patterns in a different modality, so it can generate similar encodings for both. As a result, this method naturally generates compact patient representations that are resilient to missing data. The entire process is summarized in Fig 2.

**Fig 2.**
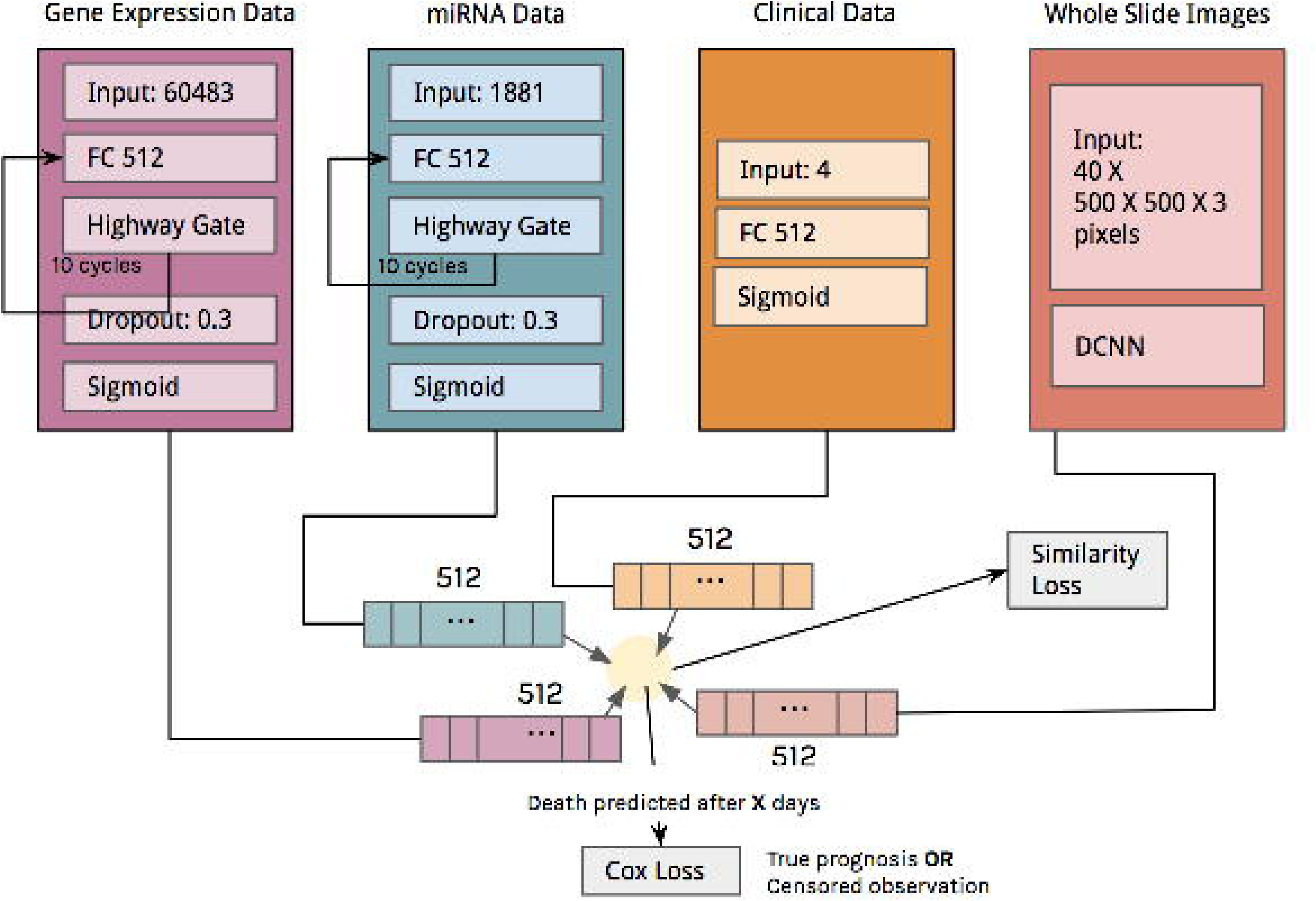
Structure of the unsupervised model: the similarity loss can be visualized as “pulling” representations of different modalities together. Each modality uses a different network architecture. For the clinical data we use fully connected layers with sigmoid activations, for the genomic data we use deep highway networks [37] and for the WSI images we use the SqueezeNet architecture [38] (see main text for architecture details). These architectures generate feature vectors that are then aggregated into a single representation and used to predict overall survival.

### Prognosis Prediction

In addition to learning strong feature representations, the model must also accurately predict prognosis. Because this is a survival data problem, we aim to maximize the concordance score or C-index. Previous research has defined the Cox loss function [39] as the best way to maximize concordance differentiably. Thus, we add a final prediction layer that maps the 512 feature vector to a survival prediction. We use the standard formulation of Cox loss to train the model. Cox loss is defined as

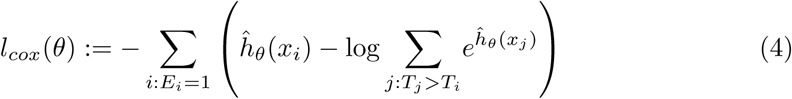

where the values *T*_*i*_, *E*_*i*_ and *x*_*i*_ are, respectively, the survival time, the censorship flag, and the data for each patient, and *ĥ*_*θ*_ represents the neural network model trained to predict survival times. The loss is computed over all patients whose lack of survival was observed. Combining with the unsupervised model, the overall loss becomes

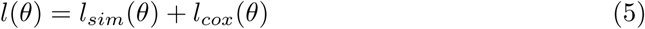

### Model Architectures

We use a dedicated CNN architecture for each data type. For the clinical data, we use fully-connected (FC) layers (Figure 2) with sigmoid activations and dropout as encoders. For the gene and microRNA data, we use highway networks as the architecture [37]. Because of the complexity and scale of WSI images, we use the CNN architecture to encode the image data. These architectures are now described in more detail.

The genomic and microRNA patient data sources are represented by dense, large one-dimensional vectors, and neural networks are not the traditional choice for such problems, for example support vector machines or random forests are more commonly used [40, 41]. However, in order to differentiably optimize the similarity and Cox loss, we must use CNNs to predict these features. Recent improvements to the state-of-art have made deep learning approaches competitive with other approaches. Thus, we use deep highway networks to train 10-layer deep feature predictors without compromising gradient flow through a neural gating approach [37]. Highway networks use LSTM-style sigmoidal gating to control gradient flow between deep layers, combating the problem of “vanishing” and “exploding” gradient in very deep feed forward neural networks (Figure 2).

In order to represent and encode WSIs, we need to develop machine learning methods that can effectively “summarize” WSIs. However, the high resolution of WSIs makes learning from them in their entirety difficult. Thus, there must be an element of stochastic sampling and filtering involved. In this work, we use a relatively simple approach to sample ROIs. We arbitrarily sample 200 (224 by 224) pixel patches at the highest resolution, then compute the “color balance” of each patch; i.e how far the average (R, G, B) color value deviates from the mean (R, G, B) value of the entire WSI (using mean-squared error). Then, we select the top 20% of these 200 patches (or 40 patches) as ROIs; this ensures that “non-representative” patches belonging to white-space and over-staining is ignored. Then, we apply a standard SqueezeNet model [38] on the 40 ROIs, with the last layer being replaced by the length-512 feature encoding predictor. The architecture is detailed in Fig 3. This model is connected to the broader network as shown in Fig 2, and is trained using the similarity and Cox loss terms. Because the SqueezeNet model is designed to be computationally efficient, we can train on a large percentage of the WSI patches without sacrificing performance.

**Fig 3.**
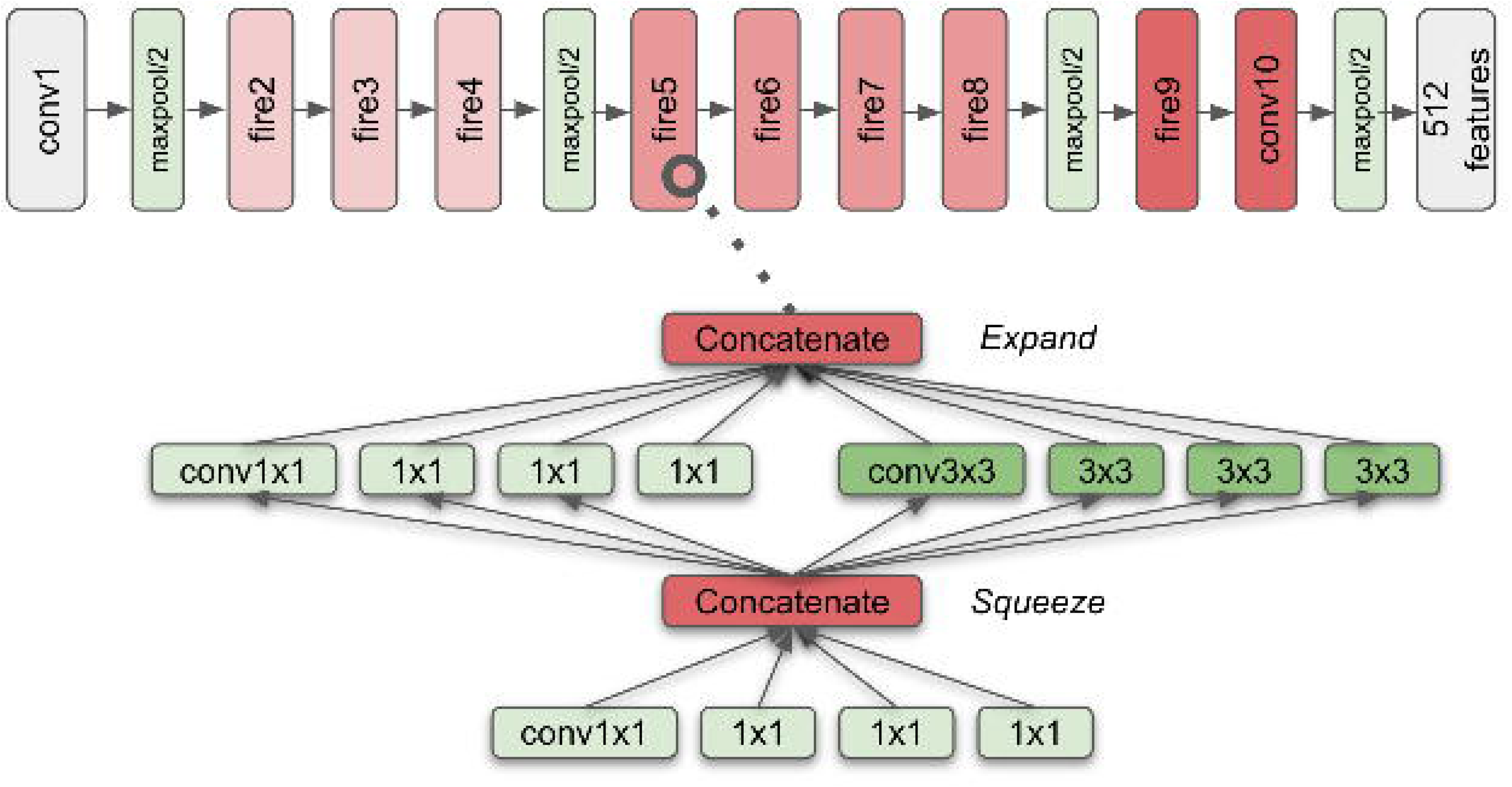
The SqueezeNet Model Architecture. The SqueezeNet architecture consists of a set of fire modules interspersed with maxpool layers. Each fire module consists of a squeeze layer (with 1×1 convolution filters) and expand layer (with a mix of 1×1 and 3×3 convolution filters). Fire module architecture helps in reducing the parameter space for faster training. We replaced the final softmax layer of the original SqueezeNet model with 512-length feature encoding predictor.

We tuned the hyperparameters of these model architectures on a validation set to find the final model parameters (Figure 2, Figure 3). To evaluate the performance of our model, we use the concordance score (C-index) on the test data set.

### Multimodal Dropout

Dropout is a commonly used regularization technique in deep neural network architectures in which some randomly selected neurons are dropped out during the training, forcing other neurons to step in to make predictions for missing neurons. This technique results in less overfitting and more generalization [42]. For multimodal dropout, instead of dropping neurons, we drop entire feature vectors corresponding to each modality, and scale up the weights of the other modalities correspondingly similar to our previous work [23]. This is applied to each data sample during training with probability *p* for each modality, to force the network to create representations that are robust to missing data modalities. We experimented with a number of different values for *P* before settling on 25% as optimal.

### Visualization

T-SNE is a commonly used visualization technique that maps points in high-dimensional vector spaces into lower-dimensions [43]. Unlike other dimensionality reduction techniques like Principal Component Analysis (PCA), T-SNE produces more visually interpretable results by converting vector similarities into joint probabilities, generating visually distinct clusters that represent patterns in the data. Here, we use T-SNE to cluster and show the relationships between our length-512 feature vectors representing patients. Because T-SNE is computationally intensive, we first used PCA to project these vectors into a 50-dimensional space, then apply T-SNE to map them into 2D space.

## Results and Discussion

### Unsupervised Learning Representations

We first evaluated the unsupervised representation learning of our model architecture by visualizing the encodings of the pancancer patient cohort (Figure 4). Clusters of patients with similar feature representations tend to have the same traits (race, sex, and cancer type), even though the model was not explicitly trained on these variables. The CNN model thus learned, in an unsupervised fashion, that factors like sex, race, and cancer type helped to identify and cluster patients across different modalities. These results suggest that the unsupervised model can effectively summarize information from multimodal data paving the way for accurate survival prediction. Perhaps even more importantly, these unsupervised encodings could act as a pancancer “patient profile”.

**Fig 4.**
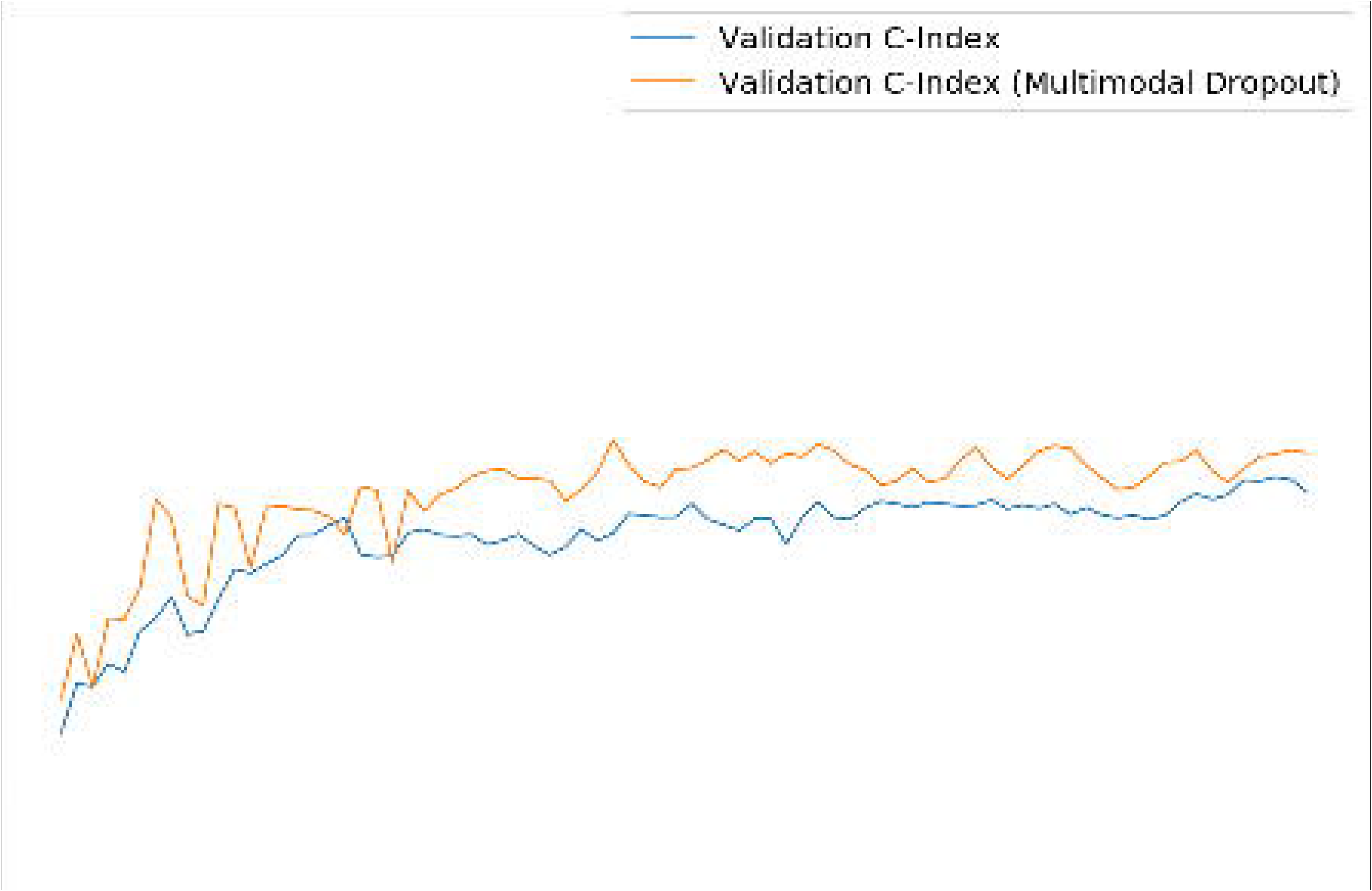
T-SNE-mapped representations of feature vectors. T-SNE-mapped representations of feature vectors for 500 patients within the testing set. The 512-length feature vectors were compressed using PCA (50 features) and T-SNE into the 2D space. These representations manage to capture relationships between patients; for example, patients with the same sex were generally clustered together (left image), and to a lesser extent, patients of the same race and same cancer type tended to be clustered as well (center and right), even when those clinical features were not provided to the model.

### Evaluation of Multimodal Dropout

Next, we evaluated the use of the multimodal dropout when integrating multi-modal clinical, gene expression, microRNA and WSIs across 20 cancer sites to predict the survival of patients. This analysis showed that the validation C-index improves when using multimodal dropout during training (Fig 5), indicating that randomly dropping-out feature vectors during training improves the network’s ability to build accurate representations from missing data. We train for 80 epochs, however the models appear to converge after 40 epochs.

**Fig 5.**
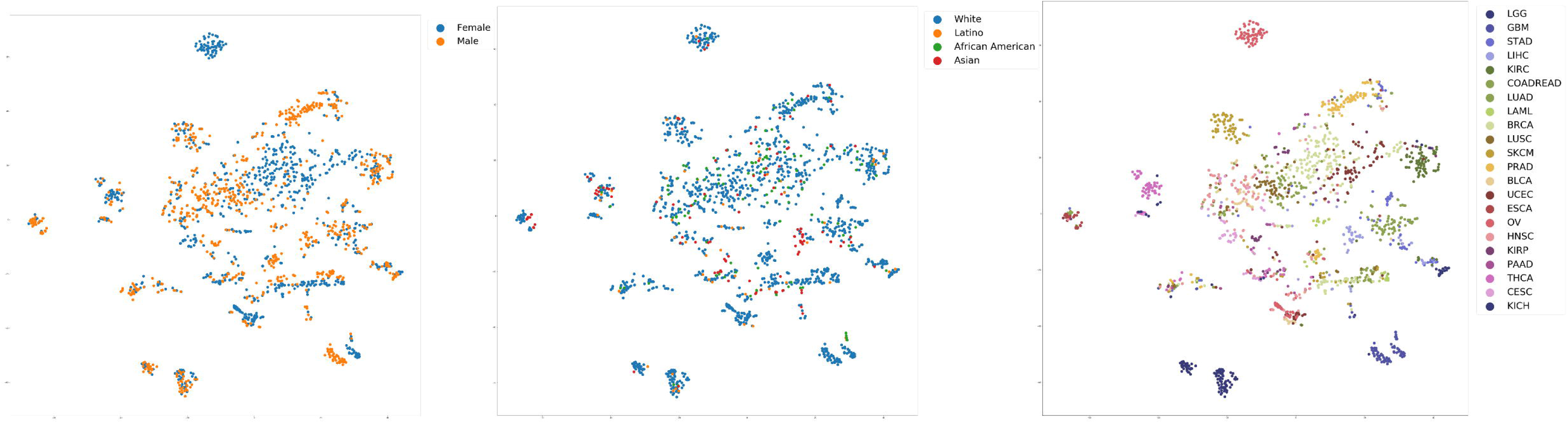
Evaluation of multimodal dropout: learning rate in terms of C-index of the model on the validation data set for predicting prognosis across 20 cancer sites combining multimodal data. The model converges after 40 epochs and shows that multimodal dropout improves the validation performance.

### Pancancer Prognosis Prediction

We then use our model on the test data set to predict prognosis for different cancer types. The model integrating clinical, mRNA, microRNA and WSI achieves an overall C-index of 0.78 on all cancers with multimodal dropout and an overall C-index of 0.75 without multimodal dropout (Table 2). For each single cancer site, multimodal dropout improves the performance by 2.7%. Models trained with multimodal dropout show an improvement (between 2-6% overall) compared to those without multimodal dropout when the same modalities are used. Note that only for mRNA, multimodal dropout did not improve the results. For the model that is trained with all modalities, many of the cancer types (14 out of 20) have a higher C-index compared to the training without multimodal dropout. Overall, prognosis prediction tended to be more accurate on cancers with more samples, but the effect of pancancer features is visible. Cancers with a low number of samples (e.g. KICH) still appear to have excellent predictive performance (i.e. best model C-index of 0.945), suggesting that training in a pancancer setting can improve the performance on underrepresented cancer sites.

**Table 2.** Model performance using C-index on the 20 studied cancer types, using different combinations of data modalities. Cancer sites are defined according to TCGA cancer codes. For each cancer the best result is bold faced. Delta refers to the relative performance of the multimodal dropout model vs. baseline. Clin=clinical data, miRNA=microRNA expression data, mRNA=gene expression data, WSI=whole slide images.

| Cancer Site | Clin + miRNA + mRNA + WSI |  |  |  | Clin + miRNA |  |  | Clin + mRNA |  |  | Clin + miRNA + mRNA |  |  | Clin + miRNA + WSI |  |  |
| --- | --- | --- | --- | --- | --- | --- | --- | --- | --- | --- | --- | --- | --- | --- | --- | --- |
|  | Single Cancer | Base-line | Multi-modal dropout | Delta | Base-line | Multi-modal dropout | Delta | Base-line | Multi-modal dropout | Delta | Base-line | Multi-modal dropout | Delta | Base-line | Multi-modal dropout | Delta |
| BLCA | 0.60 | 0.65 | 0.73 | 12.6% | 0.66 | 0.69 | 4.4% | 0.60 | 0.58 | -4.4% | 0.65 | 0.62 | -5.1% | 0.65 | 0.68 | 4.3% |
| BRCA | 0.62 | 0.77 | 0.79 | 3.0% | 0.80 | 0.80 | -0.1% | 0.57 | 0.56 | -1.9% | 0.73 | 0.73 | 0.3% | 0.77 | 0.77 | 0.0% |
| CESC | 0.52 | 0.73 | 0.76 | 4.6% | 0.77 | 0.76 | -1.2% | 0.67 | 0.62 | -6.9% | 0.74 | 0.74 | 0.4% | 0.78 | 0.76 | -2.5% |
| COADREAD | 0.58 | 0.72 | 0.74 | 3.8% | 0.78 | 0.75 | -4.8% | 0.72 | 0.58 | -20.0% | 0.77 | 0.64 | -16.9% | 0.70 | 0.74 | 4.5% |
| HNSC | 0.64 | 0.61 | 0.67 | 10.4% | 0.64 | 0.64 | 0.7% | 0.58 | 0.55 | -5.4% | 0.63 | 0.66 | 4.6% | 0.61 | 0.65 | 6.6% |
| KICH | 0.69 | 0.95 | 0.93 | -2.0% | 0.82 | 0.85 | 3.0% | 0.80 | 0.84 | 5.5% | 0.73 | 0.77 | 5.9% | 0.81 | 0.88 | 9.7% |
| KIRC | 0.78 | 0.73 | 0.73 | -0.3% | 0.70 | 0.72 | 3.1% | 0.61 | 0.65 | 5.9% | 0.65 | 0.66 | 2.7% | 0.68 | 0.61 | -11.1% |
| KIRP | 0.51 | 0.84 | 0.79 | -6.0% | 0.76 | 0.79 | 4.1% | 0.65 | 0.64 | -1.0% | 0.61 | 0.70 | 14.5% | 0.79 | 0.86 | 9.2% |
| LAML | 0.65 | 0.66 | 0.67 | 1.8% | 0.69 | 0.79 | 14.9% | 0.57 | 0.61 | 7.4% | 0.66 | 0.57 | -12.8% | 0.61 | 0.59 | -2.8% |
| LGG | 0.73 | 0.83 | 0.85 | 3.4% | 0.79 | 0.81 | 2.0% | 0.63 | 0.67 | 6.3% | 0.77 | 0.78 | 1.4% | 0.76 | 0.82 | 8.2% |
| LIHC | 0.78 | 0.72 | 0.77 | 7.6% | 0.73 | 0.74 | 2.7% | 0.64 | 0.69 | 7.7% | 0.68 | 0.67 | -1.8% | 0.70 | 0.77 | 11.2% |
| LUAD | 0.72 | 0.72 | 0.73 | 1.3% | 0.72 | 0.72 | -0.9% | 0.63 | 0.58 | -8.9% | 0.73 | 0.69 | -5.1% | 0.69 | 0.77 | 10.5% |
| LUSC | 0.63 | 0.67 | 0.66 | -0.9% | 0.72 | 0.67 | -6.5% | 0.50 | 0.51 | 2.1% | 0.62 | 0.60 | -2.9% | 0.67 | 0.68 | 0.5% |
| OV | 0.54 | 0.63 | 0.67 | 6.4% | 0.65 | 0.63 | -2.2% | 0.47 | 0.52 | 11.5% | 0.59 | 0.61 | 3.5% | 0.62 | 0.69 | 10.4% |
| PAAD | 0.57 | 0.71 | 0.74 | 3.5% | 0.68 | 0.71 | 3.8% | 0.57 | 0.61 | 7.6% | 0.59 | 0.64 | 8.9% | 0.69 | 0.69 | 0.3% |
| PRAD | 0.76 | 0.77 | 0.81 | 0.0% | 0.64 | 0.64 | 26.9% | 0.60 | 0.58 | -3.5% | 0.59 | 0.78 | 32.8% | 0.53 | 0.60 | 13.4% |
| SKCM | 0.54 | 0.68 | 0.72 | 5.2% | 0.68 | 0.68 | -0.1% | 0.56 | 0.55 | -0.1% | 0.58 | 0.72 | 24.3% | 0.67 | 0.72 | 6.8% |
| STAD | 0.60 | 0.76 | 0.78 | 2.6% | 0.75 | 0.76 | 1.5% | 0.63 | 0.54 | -13.9% | 0.80 | 0.69 | -14.1% | 0.72 | 0.74 | 2.6% |
| THCA | 0.53 | 0.95 | 0.90 | -4.8% | 0.97 | 0.95 | -2.6% | 0.82 | 0.54 | -34.2% | 0.70 | 0.83 | 18.7% | 0.93 | 0.94 | 1.4% |
| UCEC | 0.67 | 0.85 | 0.85 | 0.6% | 0.81 | 0.85 | 4.3% | 0.63 | 0.63 | 0.0% | 0.66 | 0.78 | 18.2% | 0.77 | 0.80 | 3.0% |
| Average Improvement | 2.9% |  |  |  | 1.5% |  |  | -2.4% |  |  | 5.5% |  |  | 4.3% |  |  |
| Pancancer | 0.64 | 0.75 | 0.78 | 4.5% | 0.74 | 0.78 | 4.3% | 0.60 | 0.60 | -1.2% | 0.75 | 0.78 | 3.6% | 0.76 | 0.78 | 3.2% |

### Pancancer pretraining evaluation

Next, we tested if training on pancancer data actually improved the prediction of survival across each cancer site. To test this, we compared the pancancer results with models trained on each cancer site separately for the multimodal dropout model using all data modalities (i.e. Clin + miRNA + mRNA + WSI), and compared the performance for survival prediction using exactly the same test cases for each cancer site. This showed that for all cancer sites pancancer training improves the results except for KIRC, HNSC and LIHC where a drop of 2-8% was observed (Table 2, “Single Cancer” column).

### Essential data modalities

Next, we investigated using different combinations of modalities together with clinical data, to examine if the genomic and image modalities are crucial for prognosis prediction. This shows that microRNA is the most predictive modality with a pancancer C-index of 0.78 together with clinical data compared to the weakest modality, mRNA with a C-index of 0.60” (Table 2). Next, when adding WSIs to the microRNA model the C-index is also 0.78 for a pancancer evaluation but with large differences for specific cancer sites (Table 2). In addition, the clinical-microRNA-WSI model is the best model for six cancer sites, including KIRP (C-index 0.86), OV (C-index 0.69) and LUAD (C-index 0.77) suggesting that these three data modalities are sufficient and necessary for these cancer sites prognosis determination.

All previous work on prognosis prediction using genomic and WSI data has focused on specific cancer types and data sets; thus it is difficult to exactly compare our method to previous results. Christinat *et al.*. achieved the highest C-index (0.77) thus far, on renal cancer data (TCGA-KIRC). As can be seen from the table, our method performed slightly worse (0.740) on the same type of data. But, our method heavily outperforms the only multimodality classifier (0.726 versus 0.691 C-index) on lung adenocarcinoma [24]. In general there is no “fair comparison” that can be made between this method and the previous state-of-the-art, especially because most previous papers discard patients with missing data modalities, but we train and predict with missing data included. Our methods achieve comparable or better results from previous research by resiliently handling incomplete data and predicting across 20 different cancer types.

## Conclusion

In this paper, we demonstrate a multimodal approach for predicting prognosis using clinical, genomic, and WSI data. First, we developed an unsupervised method to encode multimodal patient data into a common feature representation that is independent of data type or modality. We then illustrated that these unsupervised patient encodings are highly predictive of a wide range of useful clinical features, and that patients with similar characteristics tend to cluster together in “representation-space”. These feature representations act as an integrated multi-modal patient profile, enabling machine learning models to compare and contrast patients in a systematic fashion. Thus, these encodings could be vitally useful in a number of contexts, ranging from prognosis prediction to treatment recommendation.

We then used these feature representations to predict prognosis. On 20 TCGA cancer sites, our methods achieve the overall C-index of 0.784. Furthermore, even on cancer types that have few samples (e.g. KICH), our prognostic prediction model is able to estimate prognosis with relatively high accuracy, leveraging unsupervised features and information from other cancer types to overcome data scarcity.

Our research distinguishes itself in a number of ways. It is the first attempt to build a pancancer model of prognosis. Next, we show the use of multimodal data, novel representation learning techniques, and methods such as multimodal dropout to create models that can generalize well and predict also in the absence of one or more data modalities. More specifically, while learning unsupervised relationships between clinical, genomic and image data, our proposed CNN is forced to develop a unique, consistent representation for each patient. Finally, we propose an efficient automated WSI analysis by sampling 40 ROIs per patient representing on average 15% of the non-background space on tissue slides.

### Future Work

Although we have created an algorithm to select patches from WSI images, our results, which showed only significant improvement for select cancer sites, indicates that our method to select ROIs likely can be further improved. Refining the CNN architecture used for encoding the biopsy slides is crucial to further improve the performance. Future research, likely should focus on learning which image patches are important, rather than randomly sampling patches. Furthermore, we can use more advanced, deeper architectures and advanced data augmentation. Another intriguing possibility is using transfer learning on models designed to detect low-level cellular activity like mitoses [44]. Because of the well-established connection between mitotic proliferation and cancer, this could help focus the CNN on important cellular features. Next, integrating more diverse sources of data is another key goal. In this research, memory and computing constraints prevented us from exploring other data genomic modalities in TCGA, such as DNA methylation [45, 46] and DNA copy number data [47, 48], all of which have potentially untapped, prognostically-relevant information.

## Acknowledgments

Research reported in this publication was supported by the National Institute of Biomedical Imaging and Bioengineering of the National Institutes of Health under Award Number R01EB020527. The content is solely the responsibility of the authors and does not necessarily represent the official views of the National Institutes of Health.

